## Supplementary figures and images for "Stable sequential dynamics in prefrontal cortex represents subjective estimation of time"

### Supplementary Figure 1

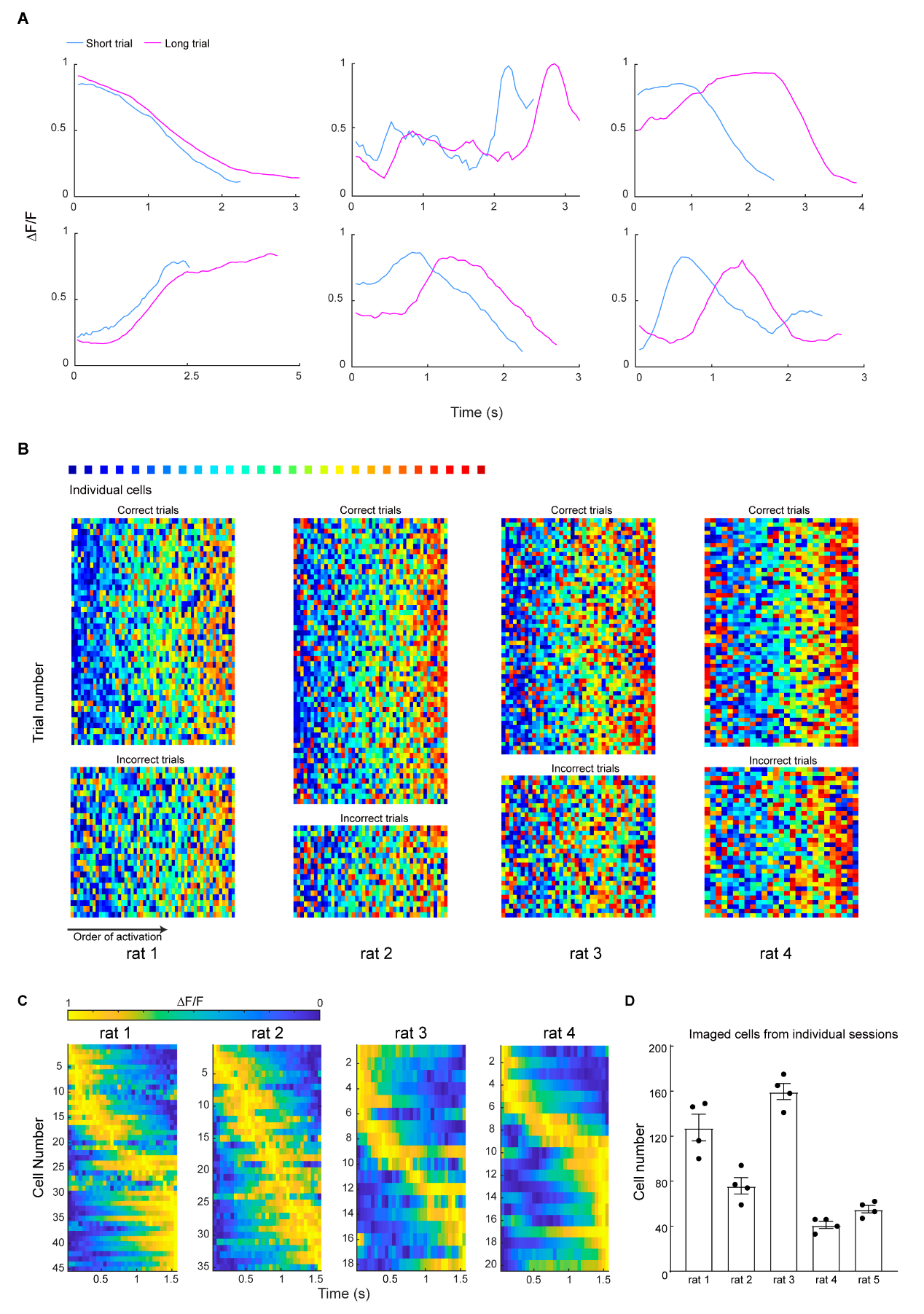

### Supplementary Figure 2

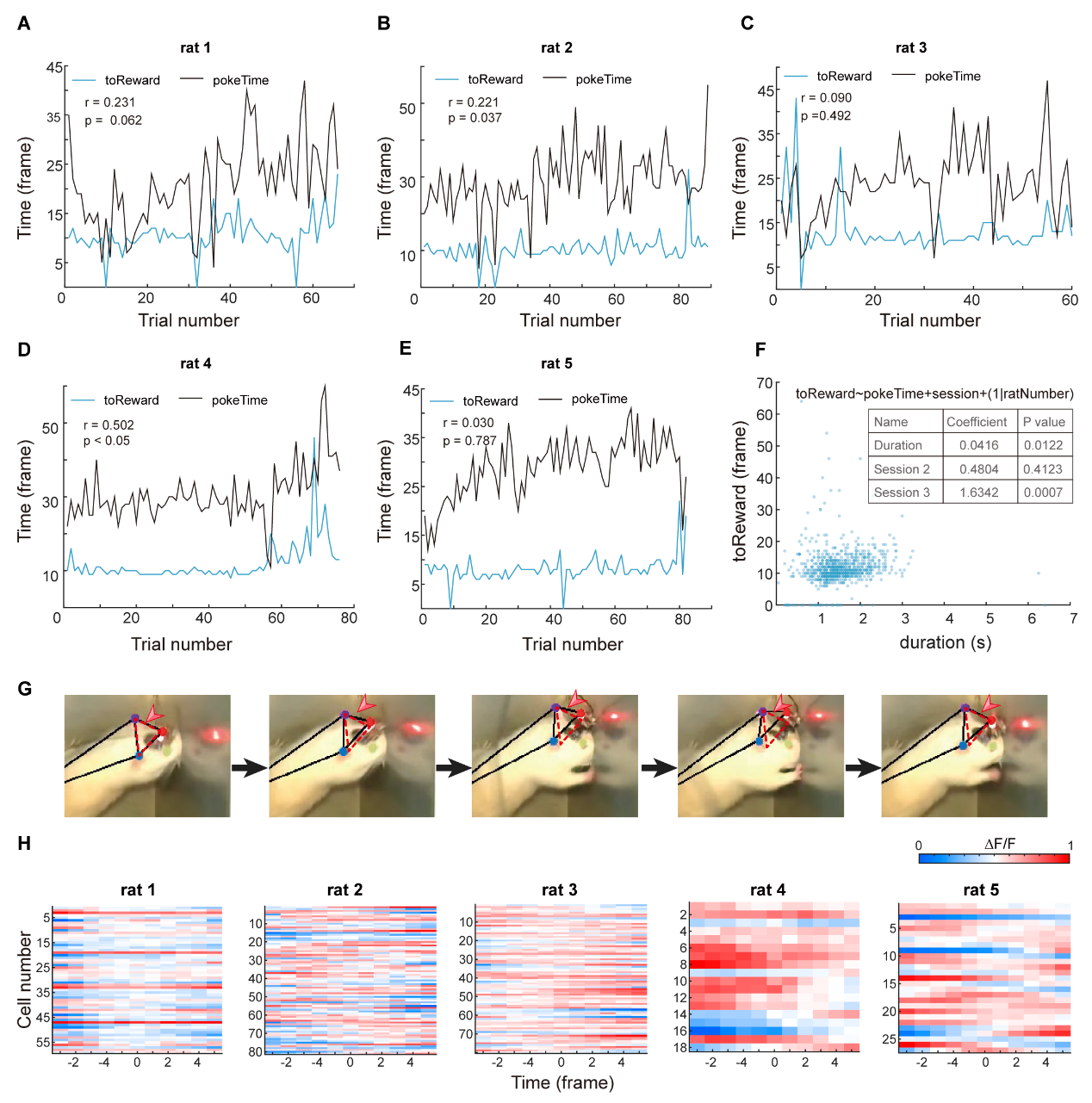

### Supplementary Figure 3

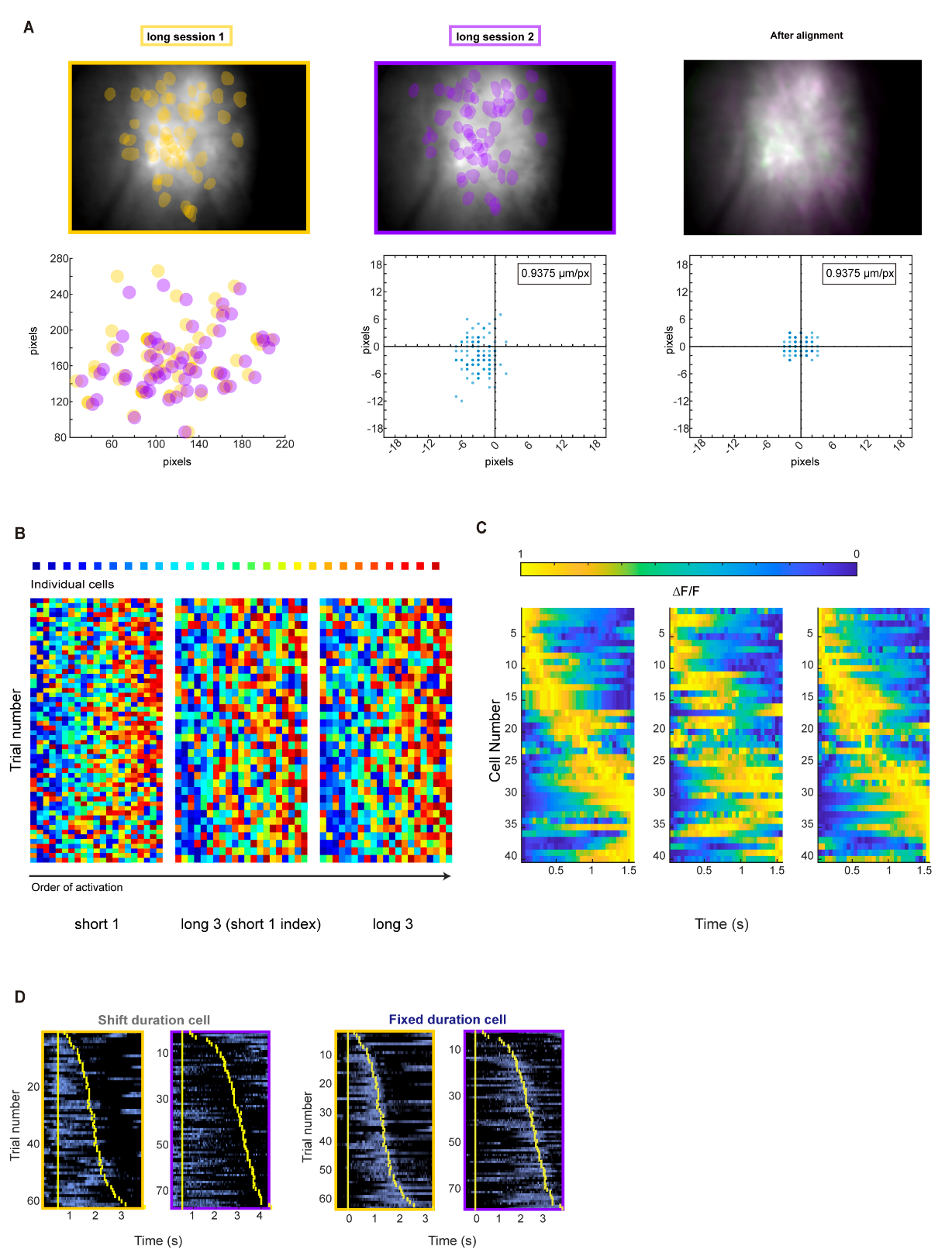

### Supplementary Figure 4

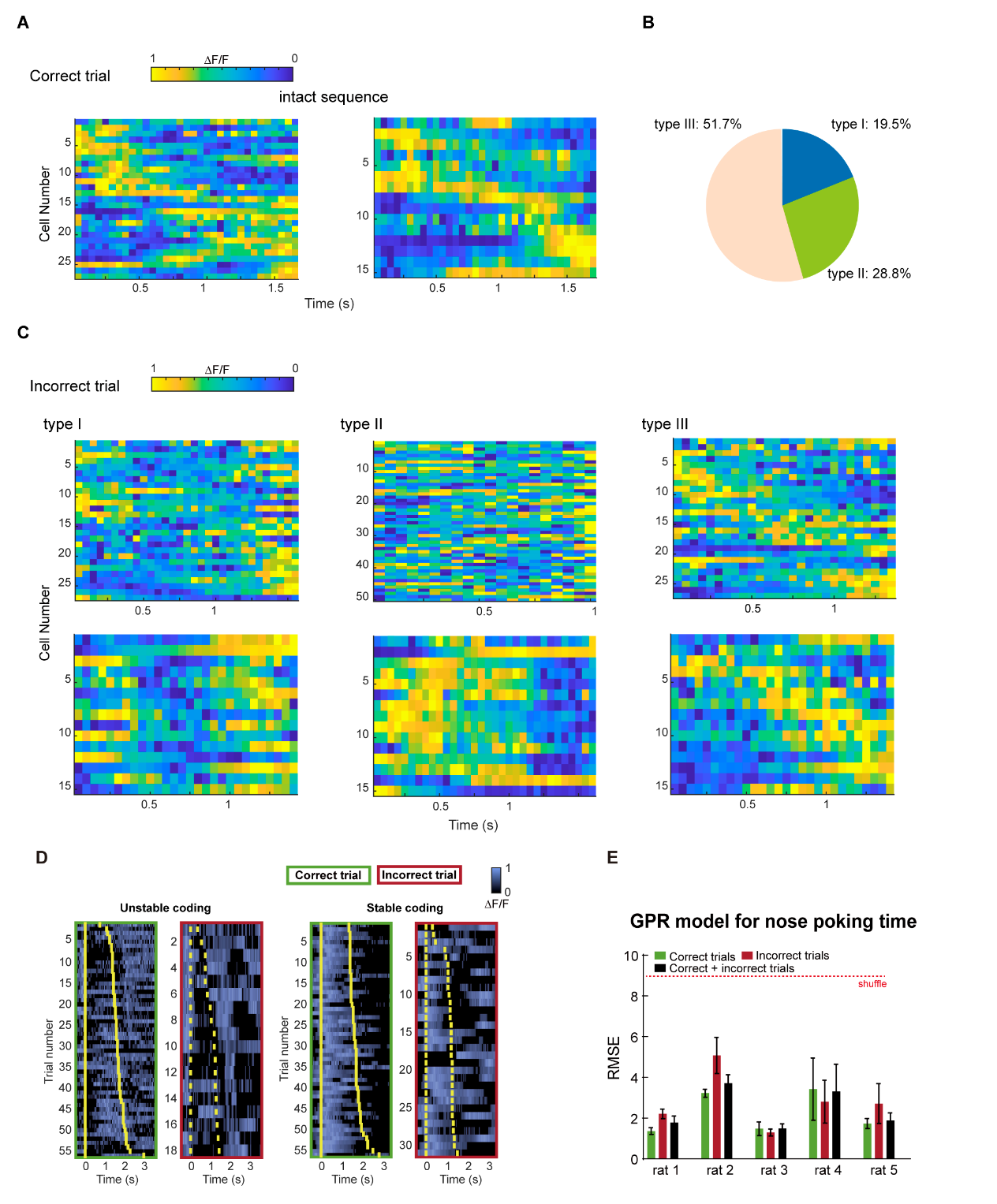

### Supplementary Figure 5

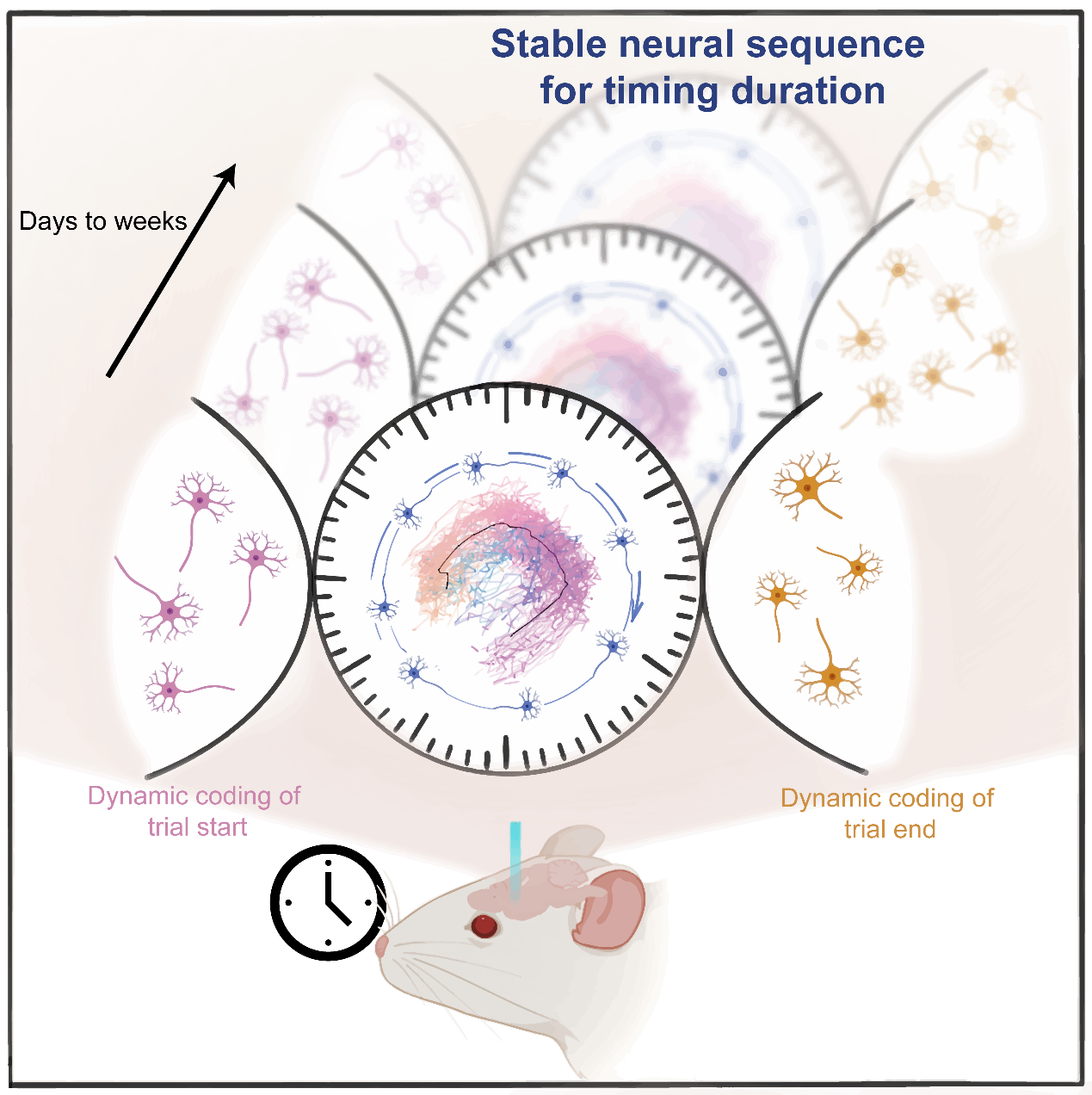
